# SMART: An open source extension of WholeBrain for iDISCO+ LSFM intact mouse brain registration and segmentation

**DOI:** 10.1101/727529

**Authors:** Michelle Jin, Joseph D. Nguyen, Sophia J. Weber, Carlos A. Mejias-Aponte, Rajtarun Madangopal, Sam A. Golden

## Abstract

Mapping immediate early gene (IEG) expression across intact brains is becoming a popular approach for identifying the brain-wide activity patterns underlying behavior. Registering whole brains to an anatomical atlas presents a technical challenge that has predominantly been tackled using automated voxel-based registration methods; however, these methods may fail when brains are damaged or only partially imaged, can be challenging to correct, and require substantial computational power. Here we present an open source package in R called SMART (<u>s</u>emi-<u>m</u>anual <u>a</u>lignment to reference templates) as an extension to the WholeBrain framework for automated segmentation and semi-automated registration of experimental images to vectorized atlas plates from the Allen Brain Institute Mouse Common Coordinate Framework (CCF).

The SMART package was created with the novice programmer in mind and introduces a streamlined pipeline for aligning, registering, and segmenting large LSFM volumetric datasets with the CCF across the anterior-posterior axis, using a simple ‘choice game’ and interactive user-friendly menus. SMART further provides the flexibility to register partial brains or discrete user-chosen experimental images across the CCF, making it compatible with analysis of traditionally sectioned coronal brain slices. In addition to SMART, we introduce a modified tissue clearing protocol based on the iDISCO+ procedure that is optimized for uniform Fos antibody labeling and tissue clearing across whole intact mouse brains. Here we demonstrate the utility of the SMART-WholeBrain pipeline, in conjunction with the modified iDISCO+ Fos procedure, by providing example datasets alongside a full user tutorial. Finally, we present a subset of these data online in an interactive web applet. The complete SMART package is available for download on GitHub.

## Introduction

Neural activity mapping using immediate early gene (IEG) expression has been invaluable for the identification of brain-wide activity patterns underlying behavior ^1, 2^. Recent advances in whole-mount tissue clearing and deep tissue immunohistochemistry (IHC) ^3-7^, coupled with improvements in volumetric fluorescent imaging ^8, 9^ and open-source brain mapping pipelines ^1, 10^ have made it possible to analyze IEG expression across intact brains. When such methods are combined with viral and transgenic tools used for cell-type and circuit specificity, novel structure-activity-function relationships at multiple spatial scales can be identified.

iDISCO ^3^ is a whole-mount clearing method that enables deep antibody diffusion and immunolabelling with a variety of antibodies that work well in traditional IHC. More recently, iDISCO+ was introduced to reduce non-isotropic tissue shrinkage and to better preserve brain morphology ^1^, permitting more reliable registration of cleared brains to established brain atlases. iDISCO+ also works well for staining Fos, an immediate-early gene marker (IEG) commonly used as a proxy for recent neural activation ^11, 12^ making iDISCO+ a valuable tool in examining relationships between brain wide activity patterns and behavior.

Several notable impediments to the widespread adoption of Fos whole brain imaging are the difficulty in properly registering images to a standardized brain atlas, segmenting the signal of interest through a full volume, and analyzing the subsequently large datasets. WholeBrain (http://wholebrainsoftware.org), a recently released open source R package ^10^, allows for semi-automated registration and automated segmentation of vectorized atlas plates from the Allen Brain Institute Mouse Common Coordinate Framework (CCF) and Reference Atlas ^13^ to experimental images. A major advantage of this approach is its scale-invariant registration process, which accommodates a wide variety of imaging resolutions. These factors, combined with the package’s minimal processing requirements, make WholeBrain an analytical registration framework that encourages widespread adoption of neural mapping through its accessibility to non-specialized labs.

However, it is difficult to register large imaging datasets in the base WholeBrain package, such as those generated by high z-step resolution scans of cleared intact mouse brains using light-sheet fluorescent microscopy (LSFM), for two main reasons. First, users are required to manually identify the anatomical coordinates along the anterior-posterior axis (AP) that correspond to the paired reference atlas plate. Second, tissue clearing methods such as iDISCO+ and CLARITY ^14^ result in non-uniform tissue morphing along the AP axis compared to the AP axis of the template atlas plates. Together, these caveats do not allow for a simple linear transformation of estimated AP coordinates between experimental images and the reference atlas, creating a significant challenge when registering an intact whole brain. Further, for datasets with a high-resolution z-step size, a potential issue is the segmentation of the same cell across multiple adjacent images as WholeBrain attempts to assign 2D segmentation to 3D space, resulting in duplicate cell counts on adjacent images.

To address these difficulties, we developed an open source software package in R called SMART (<u>s</u>emi-<u>m</u>anual <u>a</u>lignment to reference templates), built as an extension to the existing infrastructure from WholeBrain. SMART provides a streamlined pipeline for registration, segmentation and downstream analysis of large high-resolution partial and whole brain imaging datasets. Notably, SMART is built with the novice programmer in mind and requires minimal R or programming experience. It includes a console interface to enhance the user experience of WholeBrain and provides additional approaches to graphically represent and parse the mapped datasets. Lastly, SMART flexibly allows for registration of discrete, user-chosen experimental images across a mouse brain (termed partial datasets), making it a useful approach to analyze more traditional physically sectioned brain slice imaging datasets.

Here, we demonstrate the features of the SMART package in example mouse whole brain datasets that were cleared and immunolabelled for the activity marker Fos. We used a modified version of the iDISCO+ immunolabeling and clearing protocol for whole-brain Fos labelling ^1^ (we included optimization steps that enhance Fos antibody penetration and signal-to-noise ratio) and acquired volumetric image datasets at single-cell resolution using LSFM. We illustrate the utility of SMART in registering the autofluorescence channel and segmenting the Fos channel of an example dataset and demonstrate new visualization tools to graphically represent the mapped dataset. Detailed instructions for installation and use of the SMART package can be found at our tutorial webpage (https://mjin1812.github.io/SMART), while the source code can be found in our GitHub repository (https://github.com/mjin1812/SMART). In our tutorial, we also include a sample dataset containing both raw and fully analyzed data and provide a step-by-step tutorial on how to use SMART’s functions and features to implement these analyses.

With SMART, we extend the WholeBrain framework to better align datasets of non-uniformly morphed cleared brains along the anterior-posterior axis to a standardized atlas. We also present a standardized analytical pipeline addressing a key need to make mouse neural mapping projects more accessible and processing of whole brain datasets user-friendly.

## Methods

### Subjects and behavior

We used two 3-6-month-old CD-1 male mice (Charles River Labs, CD-1 IGS; strain code 022) and one Thy-1 GFP transgenic (Jackson Labs, B6;CBA-Tg(Thy1-EGFP)SJrs/NdivJ; stock 011070) mouse that we maintained on a reverse 12 h light/dark cycle (lights off at 8:00 A.M.). We performed all surgical procedures in accordance with the *Guide for the Care and Use of Laboratory Animals* (Ed 8, 2011), under protocols approved by the local Animal Care and Use Committee. CD-1 mice used for example datasets were taken from a cohort that underwent operant aggression self-administration from a control and reinforced aggression condition, as previously described^15-17^.

### Tissue clearing and immunolabeling using modified iDISCO+

We used a modified version of the iDISCO+ protocol (see http://www.idisco.info for original protocol) to achieve uniform immunostaining and tissue clearing across intact mouse brains. The timeline for the entire procedure is shown in Table 1 and a list of reagents and suggested suppliers are shown in Table 2. We also indicate time ranges where appropriate to allow for some flexibility in scheduling.

**Table 1.** A table of step-by-step instructions for our modified version of the iDISCO+ immunolabelling and tissue clearing protocol for staining c-Fos in intact mouse brains. Times recommended for each step are listed along with sample preparation details and conditions.

| Step # | Chemical | Time | Temperature | Container | Notes |
| --- | --- | --- | --- | --- | --- |
| 1a. | 1x PBS / 4% PFA | - | - | - | Transcardial perfusion |
| 1b. | 4% PFA | Overnight | 4°C | 15 mL Falcon tube |  |
| 1c. | 1x PBS | 30-60 min x 3 times | Room Temperature (RT) | 15 mL Falcon tube | Mix using a rotary shaker |
| NOTE: Prior to the next step, samples can be transferred to PBS with 0.1% Sodium Azide and stored long term at 4°C if necessary. |  |  |  |  |  |
| 2a. | 100% DI H<sub>2</sub>O | 1-h x 2 | RT | 15 mL Falcon tube | Mix using a rotary shaker |
| 2b. | 20% MeOH | 1.5-h | RT | 15 mL Falcon tube | Mix using a rotary shaker |
| 2c. | 40% MeOH | 1.5-h | RT | 15 mL Falcon tube | Mix using a rotary shaker |
| 2d. | 60% MeOH | 1.5-h | RT | 15 mL Falcon tube | Mix using a rotary shaker |
| 2e. | 80% MeOH | 1.5-h | RT | 15 mL Falcon tube | Mix using a rotary shaker |
| 2f. | 100% MeOH | 1.5-h | RT | 15 mL Falcon tube | Mix using a rotary shaker |
| 2g. | 100% MeOH | Overnight | RT | 15 mL Falcon tube | Mix using a rotary shaker |
| 3a. | 66% DCM/33% MeOH | 2-h | RT | 15 mL Falcon tube | Mix using a rotary shaker and wait till the brain sinks to the bottom |
| 3b. | 66% DCM/33% MeOH | Overnight | RT | 15 mL Falcon tube | Mix using a rotary shaker and wait till the brain sinks to the bottom |
| 3c. | 100% MeOH | 3-h | RT | 15 mL Falcon tube | Mix using a rotary shaker |
| 4a. | 100% MeOH | 3-h | RT | 15 mL Falcon tube | Mix using a rotary shaker |
| 4b. | 5% H<sub>2</sub>O<sub>2</sub> in 100% MeOH | Overnight | 4°C | 15 mL Falcon tube | Mix using a rotary shaker |
| 4c. | 80% MeOH | 1.5-h | RT | 15 mL Falcon tube | Mix using a rotary shaker |
| 4d. | 60% MeOH | 1.5-h | RT | 15 mL Falcon tube | Mix using a rotary shaker |
| 4e. | 40% MeOH | 1.5-h | RT | 15 mL Falcon tube | Mix using a rotary shaker |
| 4f. | 20% MeOH | 1.5-h | RT | 15 mL Falcon tube | Mix using a rotary shaker |
| 4g. | 100% DI H<sub>2</sub>O | 1.5-h | RT | 15 mL Falcon tube | Mix using a rotary shaker |
| 5a. | 1X PBS | 1-h | RT | 15 mL Falcon tube | Mix using a rotary shaker |
| 5b. | PTx0.5 | 1-h | 37°C | 15 mL Falcon tube | Mix using a rotary shaker |
| 5c. | PTx0.5 | 1-h | 37°C | 15 mL Falcon tube | Mix using a rotary shaker |
| 5d. | PTx0.5 | Overnight | 37°C | 15 mL Falcon tube | Mix using a rotary shaker |
| NOTE: Prior to the next step, samples can be transferred to PTx0.5 with 0.1% Sodium Azide and stored for a few days at 37°C if necessary. |  |  |  |  |  |
| 6a. | 78.6% PTx0.5/1.4% Glycine/20% DMSO | 48-h | 37°C | 15 mL Falcon tube | Mix using a rotary shaker. Incubation time can be extended by 24 to 48-h if needed. |
| 6b. | 84% PTx0.5/6% NDS/10% DMSO | 48-h | 37°C | 15 mL Falcon tube | Mix using a rotary shaker. Incubation time can be extended by 24 to 48-h if needed. |
| NOTE: For the next step, we observed that five sequential additions of 1° antibody at 1:1000 dilution helped improve penetration and labeling over a single bolus addition at a 1:200 dilution. |  |  |  |  |  |
| 7a. | 92% PTwH0.5/3% | 24-h | 37°C | 1.2 mL Corning cryovial with external threads | Mix using a rotary shaker |
|  | NDS/5% DMSO<br>+ 1:1000 1° Ab |  |  |  |  |
| 7b. | 1° Ab booster | 24-h | 37°C | 1.2 mL Corning<br>cryovial with<br>external threads | Mix using a rotary shaker |
| 7c. | 1° Ab booster | 24-h | 37°C | 1.2 mL Corning<br>cryovial with<br>external threads | Mix using a rotary shaker |
| 7d. | 1° Ab booster | 24-h | 37°C | 1.2 mL Corning<br>cryovial with<br>external threads | Mix using a rotary shaker |
| 7e. | 1° Ab booster | 72-h | 37°C | 1.2 mL Corning<br>cryovial with<br>external threads | Mix using a rotary shaker. |
| 8a. | PTwH0.5 | 24-h | 37°C | 15 mL Falcon tube | Mix using a rotary shaker |
| 8b. | PTwH0.5 | 24-h | 37°C | 15 mL Falcon tube | Mix using a rotary shaker |
| 8c. | PTwH0.5 | 24-h | 37°C | 15 mL Falcon tube | Mix using a rotary shaker |
| 8d. | PTwH0.5 | 24-h | 37°C | 15 mL Falcon tube | Mix using a rotary shaker |
| NOTE: Prior to the next step, samples can be transferred to PTwH0.5 with 0.1% Sodium Azide and stored for a few days at 37°C if necessary. For the next step, we observed that five sequential additions of 2° antibody at 1:500 dilution helped improve penetration and labeling over a single bolus addition at a 1:100 dilution. |  |  |  |  |  |
| 9a. | 97%<br>PTwH0.5/3%<br>NDS + 1:500 2°<br>Ab | 24-h | 37°C | 1.2 mL Corning<br>cryovial with<br>external threads | Mix using a rotary shaker |
| 9b. | 2° Ab booster | 24-h | 37°C | 1.2 mL Corning<br>cryovial with<br>external threads | Mix using a rotary shaker |
| 9c. | 2° Ab booster | 24-h | 37°C | 1.2 mL Corning<br>cryovial with<br>external threads | Mix using a rotary shaker |
| 9d. | 2° Ab booster | 24-h | 37°C | 1.2 mL Corning<br>cryovial with<br>external threads | Mix using a rotary shaker |
| 9e. | 2° Ab booster | 48-h | 37°C | 1.2 mL Corning<br>cryovial with<br>external threads | Mix using a rotary shaker |
| 10a. | PTwH0.5 | 24-h | 37°C | 15 mL Falcon tube | Mix using a rotary shaker |
| 10b. | PTwH0.5 | 24-h | 37°C | 15 mL Falcon tube | Mix using a rotary shaker |
| 10c. | PTwH0.5 | 24-h | 37°C | 15 mL Falcon tube | Mix using a rotary shaker |
| 10d. | PTwH0.5 | 24-h | 37°C | 15 mL Falcon tube | Mix using a rotary shaker |
| NOTE: Prior to the next step, samples can be transferred to PBS with 0.1% Sodium Azide and stored long term at 4°C if necessary. |  |  |  |  |  |
| 11a. | 100% DI H<sub>2</sub>O | 1-h | RT | 15 mL Falcon tube | Mix using a rotary shaker |
| 11b. | 20% MeOH | 1.5-h | RT | 15 mL Falcon tube | Mix using a rotary shaker |
| 11c. | 40% MeOH | 1.5-h | RT | 15 mL Falcon tube | Mix using a rotary shaker |
| 11d. | 60% MeOH | 1.5-h | RT | 15 mL Falcon tube | Mix using a rotary shaker |
| 11e. | 80% MeOH | 1.5-h | RT | 15 mL Falcon tube | Mix using a rotary shaker |
| 11f. | 100% MeOH | 1.5-h | RT | 15 mL Falcon tube | Mix using a rotary shaker |
| 12a. | 100% MeOH | 1.5-h | RT | 15 mL Falcon tube | Mix using a rotary shaker |
| 12b. | 66% DCM/33%<br>MeOH | 2-h | RT | 15 mL Falcon tube | Mix using a rotary shaker and<br>wait till the brain sinks to the<br>bottom |
| 12c. | 66% DCM/33% MeOH | 2-h | RT | 15 mL Falcon tube | Mix using a rotary shaker and wait till the brain sinks to the bottom |
| 12d. | 66% DCM/33% MeOH | Overnight | RT | 15 mL Falcon tube | Mix using a rotary shaker and wait till the brain sinks to the bottom |
| 12e. | 100% DCM | 3-h | RT | Wide mouth amber vial with PTFE lined cap | Mix using a rotary shaker and wait till the brain sinks to the bottom |
| 12f. | 100% DCM | 3-h | RT | Wide mouth amber vial with PTFE lined cap | Mix using a rotary shaker and wait till the brain sinks to the bottom |
| 13a. | DBE | 3-h | RT | Wide mouth amber vial with PTFE lined cap | Mix using a rotary shaker and wait till the brain sinks to the bottom |
| 13b. | DBE | 3-h | RT | Wide mouth amber vial with PTFE lined cap | Mix using a rotary shaker and wait till the brain sinks to the bottom |
| 13c. | DBE | 3-h | RT | Wide mouth amber vial with PTFE lined cap | Keep stationary |
| 14 | LSFM Imaging |  |  |  |  |

**Table 2.** A list of the reagents and antibodies used in our modified version of the iDISCO+ protocol for staining c-Fos in intact mouse brains. Supplier and catalogue number are listed; lot numbers are provided when possible.

| Reagent | Details | Supplier |
| --- | --- | --- |
| Phospho-c-Fos (Ser32) (D82C12) XP® Rabbit mAb | 5348S<br>(Lot 1) | Cell Signaling |
| anti-GFP/YFP (Green Fluorescent Protein (GFP) Antibody | GFP-1020<br>(Lot GFP697986) | Aves Laboratory |
| Alexa Fluor® 647 AffiniPure F(ab')<sub>2</sub> Fragment Donkey Anti-Rabbit IgG (H+L) | 711-606-152<br>(Lot # 128806) | Jackson ImmunoResearch |
| Alexa Fluor® 488 AffiniPure F(ab')<sub>2</sub> Fragment Donkey Anti-Chicken IgY (IgG) (H+L | 703-546-155<br>(Lot #127495) | Jackson ImmunoResearch |
| 10X PBS | 119-069101 | Quality Biological |
| Paraformaldehyde Prilled | 19202 | Electron Microscopy Science |
| Sodium Phosphate Monobasic Monohydrate | BDH9298 | VWR Life Science |
| Tween-20 | P9416 | Sigma Aldrich |
| Triton-X100 | X100 | Sigma Aldrich |
| Dichloromethane | 270997 | Sigma Aldrich |
| Methanol | A412SK | Fisher |
| Dimethyl Sulfoxide | D128-4 | Fisher Scientific |
| Glycine | G7126 | Sigma Aldrich |
| Heparin | H3393 | Sigma Aldrich |
| Sodium Azide | 58032 | Sigma Aldrich |
| Hydrogen Peroxide 30% | 216763 | Sigma |
| Normal Donkey Serum | 017-000-121 | Jackson ImmunoResearch |
| Benzyl ether 98% | 10801 | Sigma Aldrich |
| Falcon™ 15 mL centrifuge tubes (polystyrene) | 05-527-90 | Fisher Scientific |
| Wide Mouth Amber Vial (26 x 26mm) | CT242626-A-TEF-144 | Discount Vials |
| Corning® 1.2 mL cryogenic vials with external thread | CLS430658-500EA | Sigma Aldrich |

### Sample Collection

We anesthetized the mice with isoflurane and perfused transcardially with 200mL of 0.1 M phosphate buffered saline (PBS, pH 7.4) followed by 400mL of 4% paraformaldehyde in PBS (4% PFA, pH 7.4). We extracted brains and post-fixed them in 4% PFA (4°C, overnight). We then transferred brains to 15 mL conical polystyrene centrifuge tubes containing PBS with 0.1% Sodium Azide for long term storage at 4°C or processed them on the next day.

### Sample Pretreatment with Methanol

All sample pretreatment steps were performed in 15 mL conical tubes while gently mixing on a rotating mixer (Daigger Scientific, EF24935) and sample tubes were filled to the top with solutions to prevent oxidation. Following equilibration to room temperature (RT), we first washed brains in PBS (RT, 3 × 30 min). We then dehydrated samples using ascending concentrations of Methanol (MeOH) in deionized H_2_O (dH_2_O) – 0%, 20%, 40%, 60%, 80%, 100%, 100% MeOH (RT, 1.5 h each). Next, we performed delipidation using 66% Dichloromethane (DCM)/33% MeOH (RT, 1 × 8 h followed by overnight). We then washed samples in 100% MeOH (RT, 2 × 3 h each), prior to bleaching in a chilled H_2_O_2_ /H_2_O/MeOH solution (1 volume 30% H_2_O_2_ to 5 volumes 100% MeOH) overnight at 4°C. Next, we rehydrated samples using descending concentrations of MeOH in dH_2_O - 80%, 60%, 40%, 20%, 0% MeOH (RT, 1.5 h each). We then washed the samples first in PBS (RT, 1 × 1 h), then twice in PBS with 0.5% TritonX-100 (PTx0.5, 37°C, 2 × 1 h, followed by 1 × overnight).

### Immunolabeling

Immunolabeling was performed in 1.5 mL Nalgene cryotubes while gently mixing on the rotating mixer and samples were transferred to 15 mL conical centrifuge tubes for permeabilization, blocking and wash steps. We used a buffer containing 92% PBS/0.5% Tween-20/10ug/mL Heparin (PTwH0.5) for all wash steps and sample containers were filled to the top with solutions to prevent oxidation. We first permeabilized samples in 78.6% PTx0.5/1.4% Glycine/20% Dimethyl Sulfoxide (DMSO), and then incubated in blocking buffer containing 84% PTx0.5/6% Normal Donkey Serum (NDS)/10% DMSO (37°C, 2 d each). Next, we incubated samples in primary (1°) antibodies (anti-cFos: Phospho-c-Fos (Ser32) (D82C12) XP® Rabbit mAb, Cell Signaling Technology, #5348S Lot 1; anti-GFP, RRID:AB_10557109; Green Fluorescent Protein (GFP) Antibody, Aves Labs GFP-1020, Lot #GFP697986, RRID:AB_10000240) diluted in 92% PTwH0.5/3% NDS/5% DMSO (37°C, 7 d). We started with a 1:1000 dilution of antibody on day 1 and supplemented with 4 equal booster doses of 1° antibody (4 × 1 d) for a final dilution of 1:200 on day 5. After one week of 1° antibody treatment (5 additions over 5 days + 2 extra days), we performed washes in PTwH0.5 over 4 days (37°C, 4 × 12 h followed by 2 × 1 d). We then incubated samples in secondary (2°) antibodies (Alexa Fluor® 647 AffiniPure F(ab’)_2_ Fragment Donkey Anti-Rabbit IgG (H+L), Jackson ImmunoResearch Labs, 711-606-152, lot 128806, RRID:AB_2340625; Alexa Fluor® 488 AffiniPure F(ab’)_2_ Fragment Donkey Anti-Chicken IgY (IgG) (H+L), Jaxson ImmunoResearch Labs, 703-546-155, lot 127495, RRID:AB_2340375) diluted in 97% PTwH0.5/3% NDS (37°C, 7 d). We started with a 1:500 dilution of antibody on day 1 and supplemented with 4 equal booster doses of 2° antibody (4 × 1 d) for a final dilution of 1:100 on day 5. After one week of 2° antibody treatment (5 additions over 5 days + 2 extra days), we performed washes in PTwH0.5 over 4 days (37°C, 4 × 8-12 h followed by 2 × 1 d).

### Clearing

Initial clearing steps were performed in 15 mL conical polystyrene centrifuge tubes while gently mixing on the rotating mixer, and sample tubes were filled to the top with solutions to prevent oxidation. We first dehydrated samples using ascending concentrations of Methanol (MeOH) in deionized H_2_O (dH_2_O) – 0%, 20%, 40%, 60%, 80%, 100%, 100% MeOH (RT, 1.5 h each). We then performed delipidation using 66% Dichloromethane (DCM)/33% MeOH (RT, 2 × 3 h, followed by 1 × overnight). Next, we washed the samples in 100% DCM (RT, 2 × 3 h) to remove MeOH and then incubated them in DiBenzyl Ether (DBE) for clearing and refractive index matching (RT, 2 × 3 h, followed by 1 × overnight).

### Light-sheet fluorescent microscopy imaging

We used a light sheet microscope (UltraMicroscope II, LaVision Biotec) with an attached camera (Andor Neo sCMOS) and a 2x/0.5NA objective (MV PLAPO 2XC, Olympus) with non-corrected dipping cap. Cleared tissue was imaged coronally (olfactory bulb side up) using a customized sample platform. We took images in the 488 nm (autofluorescence or Thy1-GFP signal) and 647nm (Fos) channels and used a z-step size of 2.5 µm. We used Imspector Microscope software (v144) to control image acquisition with the following parameters: exposure = ∼100 ms, sheet NA = 0.156 (5 µm), sheet width = 80%, zoom = 0.63x, dynamic horizontal focus = 7, dynamic horizontal focus processing = blend, merge light-sheet = blend.

### Image pre-processing

We used the 3D rendering software arivis Vision 4D (3.0.0) to qualitatively check for skew in the coronal alignment in our imaging dataset. We manually corrected alignment to the coronal plane using the software’s Data Transformation Gallery and exported the images as TIFF files.

### SMART R package development

We wrote all package functions in base R ^18^. We wrote package documentation with roxygen2 ^19^. We used the magick package ^20^ to easily load, save, and modify images; to track function durations, we used the tictoc package^21^. We used the rgl ^22^, misc3D ^23^, and sunburstR ^24^ packages to generate 3D interactive plots, while we used ggplot2 to generate morph plots along the anterior-posterior axis. We used d3r ^25^ to easily convert data frames in R into JSON hierarchies. We included these packages, along with WholeBrain ^10^ (https://github.com/tractatus/wholebrain) as dependencies in SMART. We also include a dependency on the devtools package ^26^ to facilitate easy installation from Github. Finally, we used the Shiny package ^27^ to create an interactive web applet to display our example dataset (https://smartrpackage.shinyapps.io/smart_sample_dataset).

### Data storage and user reference material

Our compressed raw example image dataset is provided online at https://osf.io/y9uax/, courtesy of The Open Science Framework, a free online project management repository. It is easily extractable using 7-Zip, an open-source file archiver.

## Results

### Pipeline features

The WholeBrain package uses a scale invariant atlas by representing brain regions with nonuniform rational B-splines and categorizing them based on neuroanatomical definitions from the Allen Mouse Brain Institute Reference Atlas. Briefly, region mapping is achieved through the processes of registration, segmentation, and forward warping. During registration, the contours of the brain are segmented based on tissue autofluorescence. A set of correspondence points are generated around the tissue contour and matched with equivalent points around the contour of the atlas. Users can modify these correspondence points by adding and removing points or by changing their position within the tissue image. The correspondence points are then used as input to calculate the bending energy of the thin-plate spline deformation field. Features of interest are segmented in the image using multiresolution image decomposition; then, during the forward warp process, the deformation field is used to convert positions of segmented features into medial-lateral and dorsal-ventral atlas space. For each image, the segmentation, registration, and forward warp process must be performed, and its corresponding AP coordinate must be specified.

The SMART package builds upon the WholeBrain package to process intact whole and partial brain datasets in the following ways: 1) Users are guided through setting up parameters for the pipeline, and this information is easily fed through the registration, segmentation, and forward warp processes. These processes are automatically looped through the image dataset with simple function calls. 2) A user-friendly console interface is provided when possible. This is included during the registration process, and allows for easy addition, modification, and removal of correspondence points or reversion to the previous modification. 3) The issue of non-linear relationships between atlas AP coordinates to actual distances in the anterior-posterior axis in real imaging datasets is accounted for using semi-manual alignment to a subset of “reference” atlas templates. 4) Duplicate cell counts generated in adjacent z-planes during 2D segmentation are identified and removed, extending WholeBrain segmentation to 3D volumes. 5) Data output is saved and organized in standardized automatically generated subdirectories. 6) Additional SMART functions provide more ways to parse whole brain datasets and visualize data across regions of interest (ROIs). These features are implemented throughout or during different stages of the pipeline (Fig. 1); we explain them in depth in the remainder of this section. Notably, many of these features can be implemented on LSFM scans of partial brains or even images of single sections and are not restricted to the staining technique or image acquisition parameters used in this manuscript.

**Figure 1.**
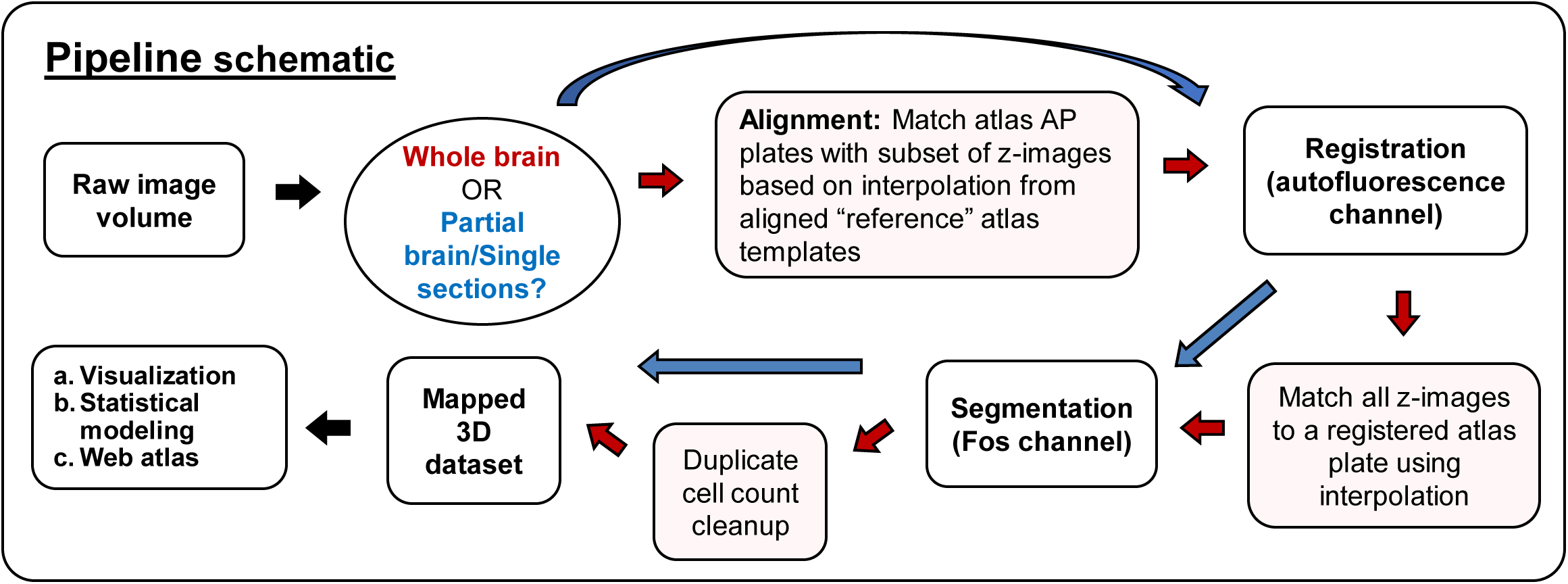
Diagram of the SMART analytical pipeline. The red arrows indicate the appropriate trajectory of an intact brain imaging dataset through steps in the pipeline, while the blue arrows indicate the appropriate trajectory for a partial brain dataset, consisting of coronal sections chosen by the user.

### Pipeline setup

Analysis parameters are organized and stored in a single variable list in R. This variable is automatically created after running an interactive function, `setup_pl()`, that guides users through entry of parameters such as the path to the desired data output folder and the z-step between images. For ease of use, the stored parameters are easily modifiable, and this variable list is the main input necessary for all remaining analytical processes in the pipeline. We include two additional functions, `im_sort()` and `get_savepaths()`, to sort image paths in the registration and segmentation channels and generate standardized subdirectories for the output of the data analysis. A comprehensive list of all SMART functions is provided in the package documentation and detailed in the online tutorial.

### Anterior-posterior alignment

SMART corrects non-linear relationships between the reference atlas and imaged tissue datasets in the AP axis by aligning a subset of reference templates to the data and interpolating the coordinates between remaining templates. The accuracy of this interpolation is evaluated using a qualitive interactive midpoint check of images between reference templates (Fig. 2, top).

**Figure 2.**
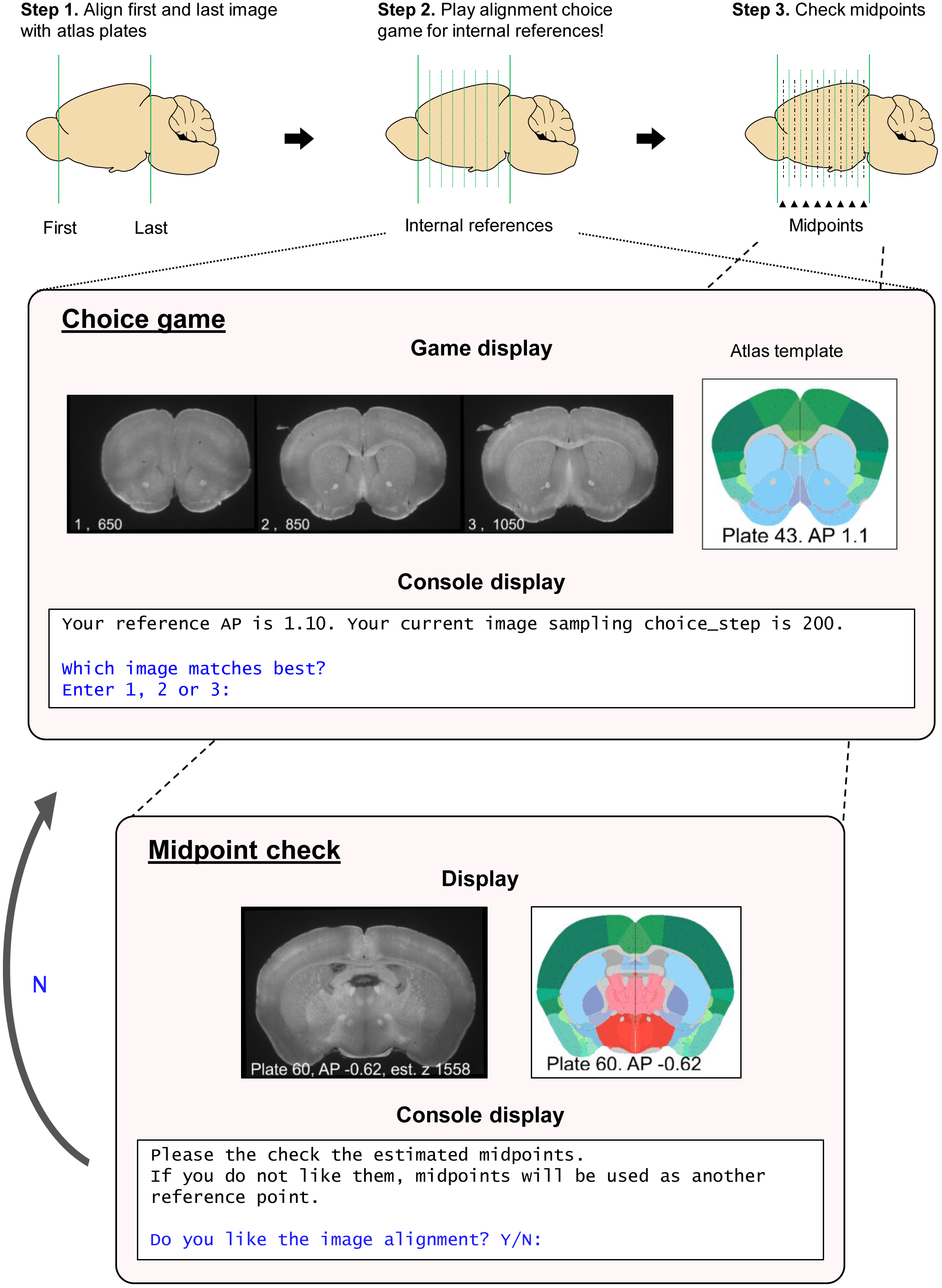
A diagram of the alignment process to reference templates. (Top) A schematic illustrating the process of qualitative alignment and inspection of midpoints of reference templates. (Middle) A visual representation of the graphical windows displayed during the choice game and the user options allowed in the R console window. The choice game is cycled through each internal reference template. During the midpoint check, the choice game is automatically played again for midpoints that are unsatisfactorily aligned; these midpoints become additional reference templates. (Bottom) A visual representation of the graphical windows displayed during the qualitative midpoint check and the user options in the R console.

During the analysis setup, only the first and last images used for analysis in the dataset need to be manually aligned with an AP coordinate. Users can do this qualitatively by comparing their images with our provided standard pdf atlas (accessible through the tutorial website) derived from the Allen Mouse Common Coordinate Framework plates used in the WholeBrain package. We suggest that users use this reference atlas, instead of other atlases such as Paxinos or Swanson, as we have found that AP coordinates and region definitions can differ across these atlas variants. Afterwards, the alignment of other images in the dataset to atlas plates within the AP range are useful for interpolation of the image set to the atlas. During the interactive setup, users specify the AP coordinates of internal reference template plates with histological landmarks that are either most familiar or most relevant to the specific experiment. If unspecified, they default to seven suggested plates (+1.91, +1.10, −0.42, −0.93, −1.94, −2.95, −3.96) across the AP axis that contain easily identifiable anatomical landmarks in the autofluorescence channel. Table 3 provides descriptions of the unique anatomical landmarks that can be consistently identified in each of these seven default coordinates plates.

**Table 3.** A list of the seven default coordinates used in the “choice game” to align experimental whole-brain mouse LSFM images to the Allen Mouse Brain Atlas. Detailed descriptions of the anatomical landmarks used to match experimental images with the plates are listed. Region acronyms used in the Allen Mouse Brain Atlas (AMBA) are also provided so users can cross validate with the AMBA (http://atlas.brain-map.org/atlas?atlas=1).

| AP coordinate | Plate number | Anatomical landmarks and characteristics |
| --- | --- | --- |
| +1.91 | 35 | 1) The folds separating the piriform area (PIR) from the orbital area (ORB) span across approximately half the horizontal length of a hemisphere. 2) There is a distinct teardrop shape to the anterior commissure, olfactory limb (aco). 3) The nucleus accumbens (ACB) has not yet appeared. |
| +1.10 | 43 | 1) The two “wings” of the corpus callosum, anterior forceps (fa) are just about to touch. 2) The diagonal band nucleus (NDB) is clearly present and extends halfway the length from the ventral-most point of the midline to the corpus callosum. |
| -0.42 | 58 | 1) The columns of the fornix (fx) and the stria medullaris (sm) have just separated; the stria medullaris is still connected in one piece. 2) The optic chiasm (och) is directly below the hypothalamus in one piece and has not yet split into two optic tracts (opt). |
| -0.93 | 63 | 1) The CA3 stratum oriens (CA3so) of the hippocampus is just beginning to show within the fimbria (fi); however, multiple layers of the CA3 aren't showing yet. 2) The columns of the fornix (fx) have migrated below the top of the third ventricle. 3) The optic tracts have migrated close to the border of the hypothalamus and amygdala. |
| -1.94 | 73 | 1) The dentate gyrus (DG) is clearly present and spans approximately 2/3rds the horizontal length of one wing of the hippocampus. 2) The ventral-most point of the granule layer of the dentate gyrus (DG-sg) is roughly the same height as the ventral-most point of the pyramidal layer of CA3 (CA3sp). 3) The fasciculus retroflexus (fr), mammillothalamic tract (mtt), and columns of the fornix (fx) are clearly seen in their appropriate positions and spaced evenly apart; the mtt is slightly closer to the fx than the fr. 4) The median eminence is (ME) is clearly visible at the bottom of the midline. |
| -2.95 | 83 | 1) The medial mamillary nucleus (MM) of the hypothalamus has receded and is connected by a thinned layer of hypothalamic tissue. 2) the posterior commissure (pc) and cerebral aqueduct have migrated down from the 3<sup>rd</sup> ventricle (V3), and they are separated from V3 by sublayers c and b (SCig-c, S-Cig-b) of the superior colliculus. |
| -3.96 | 93 | 1) The tegmental reticular nucleus (TRN) and pontine gray (PG) are easily identified and separated by the corticospinal tract (cst) and medial lemniscus (ml). 2) The brachium of the inferior colliculus (bic) protrudes out to the left and right of the midbrain, slightly beyond the middle cerebral peduncle (mcp). |

These internal reference templates are aligned to the imaging data through an interactive “choice game” where three image options are graphically presented alongside the reference atlas template (Fig 2, middle). The center image is estimated based on interpolation of the first and last aligned image, while the left and right images are further anterior and posterior options, respectively. The choice presentations become progressively closer to each other when the user chooses the middle image, while a choice of the right or left image sets them as the new middle image during the next choice cycle. The default progression of z-step choices between consecutively presented images for each coordinate is 200, 100, 30, and 10 images, though this progression user-modifiable.

Once all the internal reference coordinates have been aligned, users can run a midpoint check (Fig 2, bottom) where the midpoint atlas templates are presented alongside the estimated midpoint image from the dataset. If the alignment is unsatisfactory for a given midpoint, the user can designate them as another internal reference point and play the choice game again to improve alignment. The remaining intervening AP coordinates are estimated with interpolation.

### Interactive registration improvement

In the base WholeBrain package, removing, changing, or adding correspondence points requires a separate function call each time, and it is not possible to seamlessly revert to a previous set of correspondence points. This can be time consuming and confusing for new programmers. We incorporated all these original features into a single function call, `regi_loop()`, that provides an interactive console interface, adds the option of reverting back to the previous registration if a new modification is unsatisfactory, and automatically loops through all atlas plates that need to be registered (Fig. 3A). Further, the output of each successful atlas registration is organized and stored into an automatically generated variable list. These new features simplify the registration modification process so users can quickly improve the accuracy of their registrations (Fig. 3B). Should a registration be deemed inaccurate at a later timepoint, users can specify the specific atlas plate they want to manually correct. We also incorporate an automatic looping feature so users can obtain the initial automated registrations of all atlas plates prior to manual correction, which is useful as a first-pass quality control check for the dataset.

**Figure 3.**
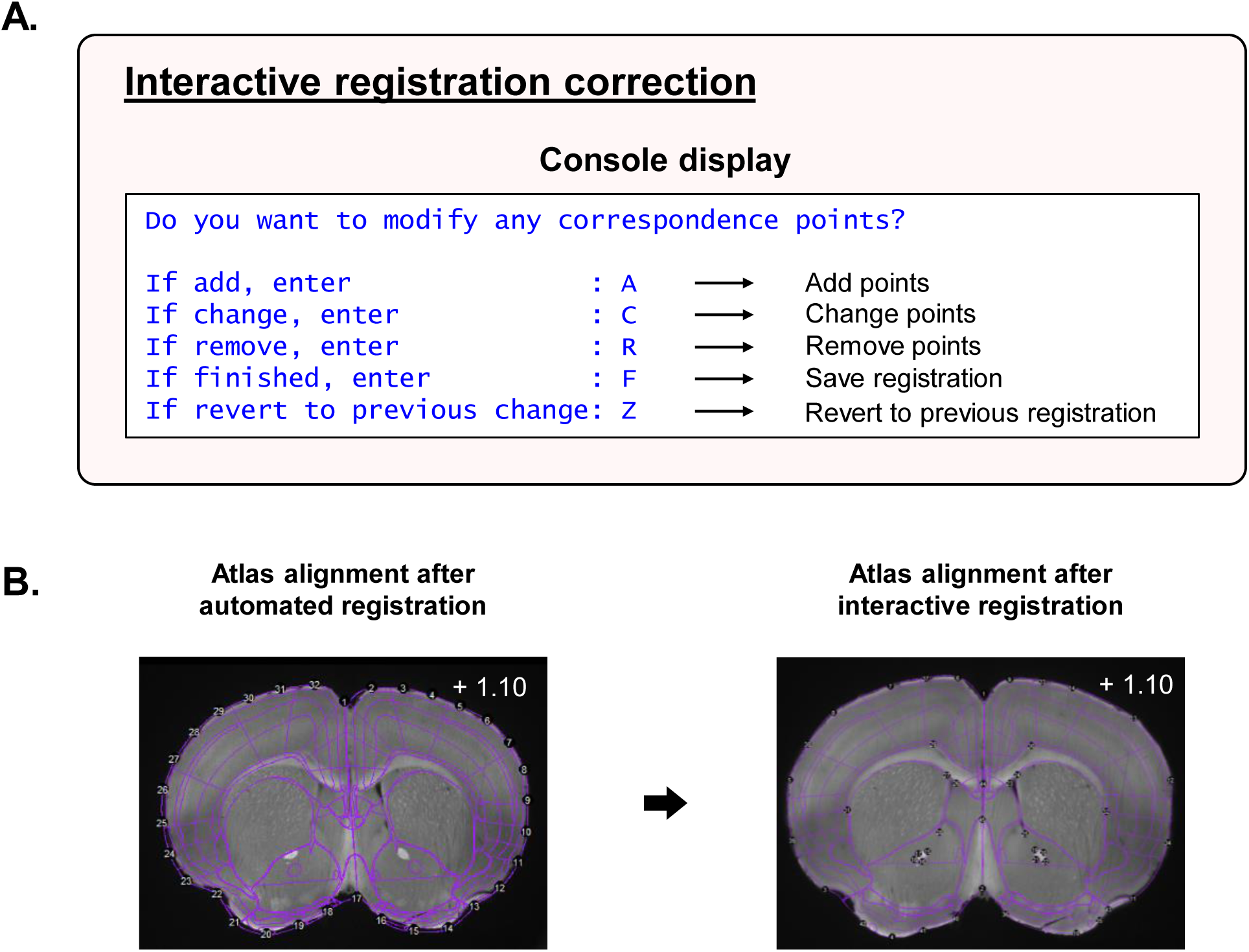
The interactive registration correction process. **(A)** A visual representation of user options in the R console display during manual correction of registrations. **(B)** An example image showing the initial registration of the atlas template to the tissue based on correspondence points around the contours of the tissue in the autofluorescence channel (left). The atlas-tissue alignment is improved following interactive registration correction (right).

### Automated segmentation, duplicate cell count cleanup, & forward warping

The features of interest segmented by WholeBrain are defined by parameters assigned to a segmentation filter. Using the filter as an additional input, SMART automatically loops through the entire image dataset and segments according to the filter parameters. For datasets with a high-resolution z-step, a major issue is the segmentation of the same cell across multiple adjacent images. To address the issue of artificially inflated cell counts, we incorporated a duplicate-cell count cleaning function, `clean_duplicates()`, in the pipeline based on user-defined distance thresholds in the z-axis and the x-y axes. The central position of the cell body is marked in the z-position containing the maximum intensity values while duplicate counts across other images are erased. The registration and corrected segmentation data, along with the setup analysis parameters are subsequently used as inputs during SMART’s forward warping function, which loops the base WholeBrain package’s warping function through the entire dataset and automatically matches each image in the segmentation with the closest registered atlas plate. The forward warp function also omits any segmented features outside the registered atlas boundaries.

We note that for brain datasets containing especially high-resolution (≥ 10x) tiled full slice images, WholeBrain’s segmentation function fails during runtime. In such cases, we suggest that users utilize other image segmentation procedures (i.e., Fiji ImageJ) to create a binary mask of the segmented cells, and then combine these binarized images with the original segmented images to retain cell-count intensity information. These new masked images can be used as the segmentation channel in the SMART pipeline.

### Pipeline validation with intact cleared brain LSFM dataset

We first validated our modified iDISCO+ clearing approach (Fig. 4A) and immunolabelled the immediate-early-gene Fos throughout the intact tissue of a Thy-1 GFP transgenic mouse (Fig. 4B); we obtained high-resolution imaging quality using our light-sheet fluorescent microscope parameters.

**Figure 4.**
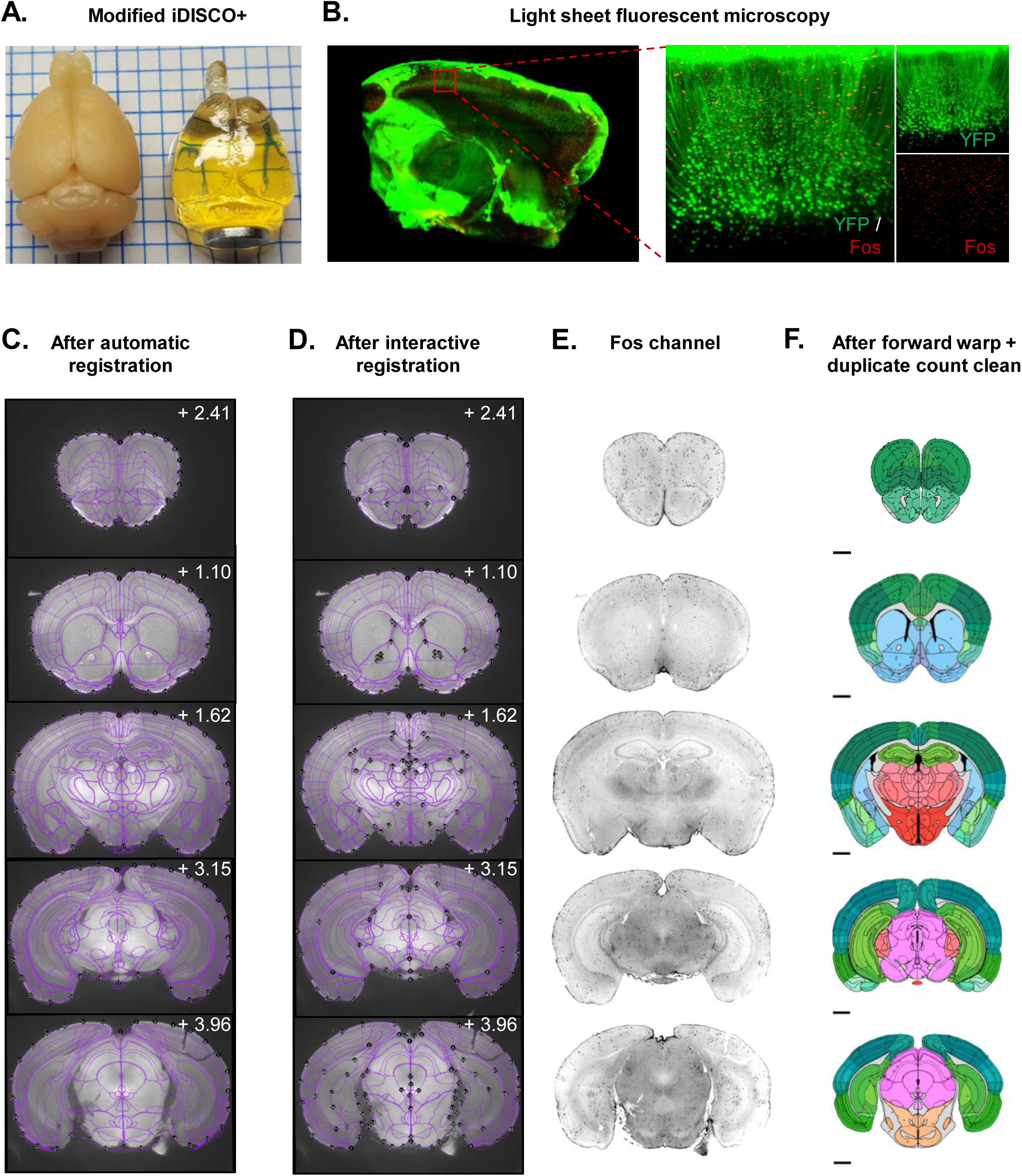
Representative images of registration correction and Fos segmentation from an example dataset. **(A)** Transparency of intact brain samples pre- and post-iDISCO+ immunolabeling and clearing. **(B)** Example LSFM tissue section of an intact cleared brain from a Thy-1 GFP transgenic mouse (left); an enlarged cortical image in the YFP and Fos imaging channels (right). **(C)** Initial atlas-tissue registrations across five representative images in the autofluorescence channel (488 nm) of the example dataset from Mouse 1 at various anterior-posterior coordinates. **(D)** The atlas-tissue registrations following interactive registration correction. **(E)** The corresponding images in the Fos channel (561 nm). Image colors were inverted in ImageJ for easier visualization. **(F)** Schematic plots showing segmented cells from **E** following cleanup of duplicate cell counts and forward warping back onto atlas space.

We then cleared and immunolabelled Fos (Fig. 4E) in a whole brain extracted from a male outbred CD-1 mouse that was perfused directly from the homecage (Mouse 1). We fed the generated LSFM dataset through the SMART pipeline, playing the choice game and manually correcting registration misalignments in the autofluorescence channel (488 nm) across all atlas plates (Fig. 4C-D). We segmented cell bodies in the Fos channel (647 nm); we then corrected the segmentation data of duplicate cell counts and performed the forward warp of all cell counts onto atlas space (Fig. 4F). We developed an interactive web applet (https://smartrpackage.shinyapps.io/smart_sample_dataset) to visualize the entirety of our pipeline output and provide a template for sharing volumetric data following analysis in SMART.

### Data organization and visualization

Extending the functionality of current data parsing features in WholeBrain, we provide the simple functions, `get_table()`, to create a table displaying region acronyms, region cell counts, and region cell count percentages, and `get_rois()`, to extract only regions of interest (ROIs) from the main dataset. The `get_rois()`function only requires users to specify region acronyms from the Allen Mouse Brain Institute Reference Atlas. SMART users can combine this capability with existing or modified visualization functions from WholeBrain to generate interactive 3D renderings and region cell count plots of user-specified ROIs. We demonstrate these features in our example dataset by generating a 3D rendering of various limbic regions (Fig. 5A) and a region cell count plot of the PFC (Fig. 5B, IL and PL). We also demonstrate two new data visualization features with our example dataset: a normalized morph plot along the anterior-posterior axis (Fig. 5C) and an interactive sunburst plot showing region cell counts and hierarchical structural relationships (Fig. 5D). We processed an additional LSFM dataset (Mouse 2) from a male outbred CD-1 mouse, exposed to operant aggression self-administration, using the SMART pipeline and illustrate the resulting brain morph (Fig. 5E) and sunburst plots (Fig. 5F) for qualitative evaluation alongside Mouse 1.

**Figure 5.**
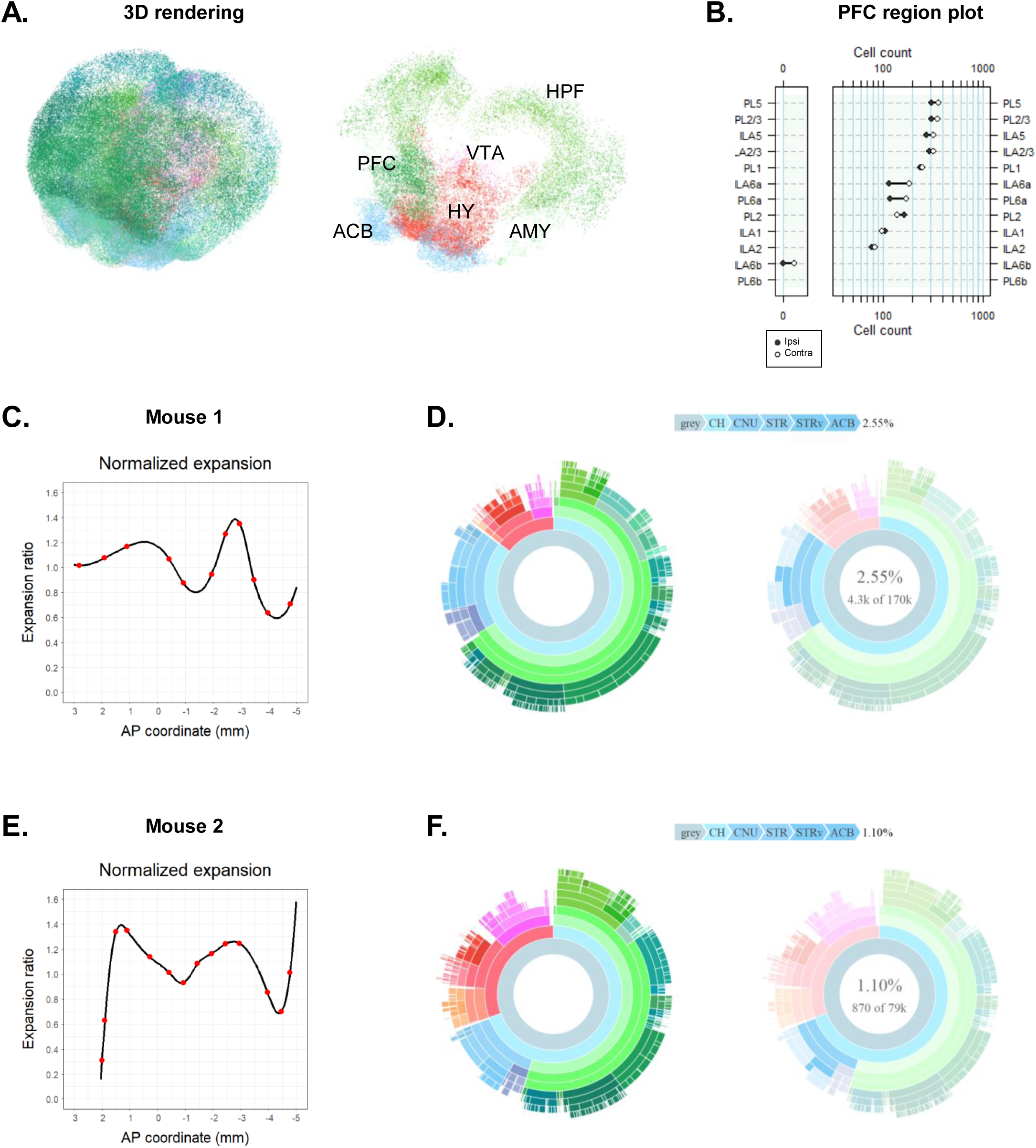
Graphical representations of mapped datasets. Figures A-D Are from the same dataset (Mouse 1) used to generate figure 4. **(A)** A 3D rendering of the entire mapped dataset (left) and a 3D rendering of the indicated regions of interest (right). **(B)** A region plot quantifying segmented cell bodies in the PFC (PL and IL). **(C)** A plot of normalized brain morph ratio across the anterior-posterior axis. The red dots indicate the positions of the aligned reference templates. The thick black line indicates the interpolated morph ratio for AP positions between reference templates. **(D)** A sunburst plot showing region cell counts and hierarchical structural relationships (left). The innermost ring represents all cell counts, with each subsequent outer ring representing child structures of the adjacent inner ring. Regions colors are based on those in the Allen Mouse Brain Atlas and arc length is proportional to overall cell counts. The plot is interactive and hovering over an arc reveals region identity, hierarchical path, and region cell count (right). **(E-F)** A brain morph plot and sunburst graph for an additional mapped dataset (Mouse 2) that performed operant aggression self-administration. <u>Abbreviations:</u> nucleus accumbens (ACB), amygdala (AMY), hippocampal formation (HPF), hypothalamus (HY), infralimbic cortex (IL), prefrontal cortex (PFC), prelimbic cortex (PL), ventral tegmental area (VTA).

## Discussion

We have developed an open source package in R which streamlines the user experience of WholeBrain and provides a built-in analytical pipeline for intact mouse brain activity mapping. SMART was designed with the novice programmer in mind. We recognized that installation of the required software and dependencies could be a major technical hurdle to using this pipeline and have provided detailed installation and setup instructions to overcome them. Users are also guided through the analysis setup process and presented with an intuitive console interface when possible. The modified registration, segmentation, and forward warp functions automatically loop through the entire imaging dataset, a valuable feature when analyzing large neuroimaging datasets. SMART also automatically creates and saves outputs in standardized data subdirectories, facilitating organized storage and sharing of datasets. Most notably, we account for the issues of non-uniform morphing and duplicate cell counts along the anterior-posterior axis in cleared tissue datasets. We demonstrated these features in an example mouse LSFM dataset generated from a brain cleared and immunolabeled for Fos using a modified iDISCO+ protocol.

A main consideration when using our pipeline is the speed of analysis. The analysis bottleneck of the base WholeBrain package is the manual correction of registered atlas plates. This remains the most time-intensive step in SMART; however, users can save significant time registering an entire brain using the intuitive interactive console interface. Using a comparable dataset to the one presented, we estimate that a well-trained user of the SMART pipeline can accurately register an entire mouse brain following three to four days of dedicated registration. The remaining segmentation, duplicate cell count cleaning, and forward warping processes can be completed within one day. A notable feature of SMART is that it allows users to save intermediate stages of analysis and return to them when convenient. This makes it possible for multiple users to work on the same data in tandem and spread the time commitment across entire teams.

The main determinant of the accuracy of region cell count mapping is the pipeline user’s own anatomical knowledge. This remains both an advantage and disadvantage of the SMART-WholeBrain approach compared to other typically intensity-based voxel registration approaches: users can easily evaluate registration quality and correct for misalignments, but exact reproducibility of registrations between different users is relative. However, we propose that using these approaches in parallel provides the best outcome. An initial registration pass using automated voxel-based approaches like ClearMap^1^, followed by SMART on brains that poorly register due to damage or partial brain datasets, provides a complete registration pipeline with qualitative checks and simple visualization of results.

With the above considered, the SMART-WholeBrain analysis framework is timely, as mapping of cell distributions across intact brain tissue is becoming an integral approach towards understanding the activity and function of neural circuits at the level of the connectome. The resurgence in interest in tissue clearing techniques within the last decade has driven new advancements in high resolution volumetric imaging methodologies, such as serial two-photon tomography ^28^ and light sheet fluorescence microscopy ^29^. Such methods faithfully capture tissue at the resolution of a single-cell or higher, but in doing so they generate unwieldy sizes in neuroimaging datasets.

Until now, there have been few streamlined attempts towards analyzing such datasets, as most approaches toward cell or axonal distribution mapping involve time-intensive manual annotation or semi-automated analysis using specialized software ^30-34^. Automated approaches towards registration of an imaging dataset to a standardized atlas are often specifically tuned to the imaging setup and to the experimental datasets being evaluated ^35, 36^ and are not readily available for others to use. Furthermore, such automated methods are generally developed from intensity-based registration software ^37^ and do not easily allow for user evaluation of registration quality or manual registration correction.

The variability and specialization of these approaches underscore the need for open source software options that are easily accessible, user-friendly, and encourage whole-brain activity mapping projects that require little specialized knowledge. Though there are other freely available region-based cell quantification pipelines with demonstrated utility ^1^, WholeBrain’s computational framework is versatile due to its minimal processing requirements, scale-invariant registration, and manual registration correction capabilities. We expect the SMART package will enhance the accessibility of the base WholeBrain package and encourage more users to undertake neural mapping projects, whether using full brain or partial brain datasets.

To support this endeavor, our example dataset is freely available and downloadable to test. We have created a website to walk users through the package installation process in R and through a tutorial of SMART’s functions using the example dataset. We also present our pipeline output in a convenient web applet to visualize registration, segmentation and other data. A more detailed package description is available at https://mjin1812.github.io/SMART/.

## Author contributions

MJ, RM, and SAG conceived of the pipeline and contributed intellectually to its development. MJ programmed, packaged, and created SMART and its supporting webpage. RM and CAMA developed the modified tissue clearing protocol. RM, CAMA and SAG conducted all imaging. MJ, SJW and JDN analyzed the dataset. JDN developed the Shiny applet used to display the example dataset. MJ, RM, and SAG wrote the manuscript with input from the other authors.

## Acknowledgments

The authors would like to thank Yavin Shaham and Bruce Hope and their lab members for their help with this project. We would like to thank Lauren E. Komer and Vadim Kashtelyan for their technical assistance. We also thank the NIDA Histology Core and staff for their support.

